# Potential biomarker of human papillomavirus 16 L1 methylation for prediction of anal intraepithelial neoplasia in men who have sex with men (MSM)

**DOI:** 10.1101/2021.01.07.425707

**Authors:** Arkom Chaiwongkot, Nittaya Phanuphak, Tippawan Pankam, Parvapan Bhattarakosol

**Author notes:** Corresponding author: Arkom Chaiwongkot, Ph.D. Division of Virology, Department of Microbiology, Faculty of Medicine, Chulalongkorn University, 1873 Rama 4 Road, Patumwan, Bangkok 10330, Thailand.

## Abstract

Quantitative measurement of human papillomavirus (HPV) 16 early promoter and L1 genes methylation were analyzed in anal cells collected from men who have sex with men (MSM) to determine the potential biomarker for screening of HPV related anal cancer. The methylation patterns of the HPV16 genes including early promoter (CpG 31, 37, 43, 52 and 58) and L1 gene (CpG 5600, 5606, 5609, 5615, 7136 and 7145) were analyzed in 178 anal samples with histology diagnosed as normal, anal intraepithelial neoplasia (AIN) 1, AIN2 and AIN3 by pyrosequencing assay. Low methylation levels of early promoter (<10%) and L1 genes (<20%) were found in all detected normal anal cells, while medium to high methylation (>20-60%) in early promoter was found 1.5% (1/67) and 5% (2/40) in AIN1 and AIN2-3 samples, respectively. Interestingly, slightly increased L1 gene methylation level (>20-60%) especially at HPV16 5’L1 regions CpGs 5600 and 5609 from normal to AIN3 were demonstrated. Moreover, negative correlation between high HPV16 L1 gene methylation at CpGs 5600, 5609, 5615 and 7145 and low percentage of CD4+ was found in AIN3 HIV positive cases. When compared methylation status of AIN2-3 to those of the normal/AIN1 lesion, the results indicated potential of using HPV16 L1 gene methylation as a biomarker for HPV related cancer screening.

## Introduction

Anal carcinoma is a rare disease found in men and women population globally with average incidence of <1-2 per 100,000[1]. However, the incidence of anal cancer was high in HIV infected women and HIV-infected men who have sex with men (MSM), accounting for 30/100,000 and 131/100,000, respectively [2]. There is association between human papillomavirus (HPV) infection and anal carcinoma in which HPV DNA was found in men (68.7-91.2%) and women (90.4-90.9%) with anal carcinoma worldwide [3]. Studies showed that HPV 16 was predominantly detected in men and women, accounting for 70-71.6% and 74-83.4%, respectively [2, 4, 5].

High prevalence of HPV infection was reported in anal cells collected particularly from HIV infected MSM. Worldwide HPV prevalences in anal cells of HIV infected and HIV uninfected MSM were 92.6% and 63.9%, respectively and HPV16 was found 35.4% and 12.5% in general HIV infected and HIV uninfected MSM, respectively [6]. Recent study in China reported high prevalence of HPV infection among HIV-infected MSM (82.69%) compared with HIV uninfected MSM (62.81%)[7]. In Bangkok, Thailand, anal HPV infection was found 85% in HIV infected MSM compared with 58.5% in HIV infected MSM in which HPV16 was detected 22.5% and 9.8% in HIV infected and HIV uninfected MSM, respectively [8]. The prevelance of anal HPV infection in northern Thailand was 80% among MSM, in which 100% and 70% were found in HIV infected MSM and HIV uninfected MSM, respectively, HPV16 was the most common high risk types, accounting for 40% in HIV infected MSM and 22% in HIV uninfected MSM [9]. It was reported that HPV16 was the most persistent high risk HPV types [10] and less likely to have spontaneous regression from cervical intraepithelial neoplasia (CIN)2-3 to normal when compared to other HPV types [11, 12]. The study in HIV uninfected MSM revealed that HPV16 showed the longest duration of infection with the lowest rate of viral clearance when compared to low risk HPV [13].

High risk HPV is considered to be the causative agent of cervical cancer and other HPV related cancers such as vulva, head and neck cancer and anal. Viral oncoproteins E6 and E7 disrupt normal function of host proteins involved in cell cycle regulation such as E6 causes p53 degradation and E7 inactivates retinoblastoma protein[14, 15]. However, the development of HPV related cancers takes more than 10-20 years, while the majority of HPV infected population has spontaneous regression [16, 17]. Up-regulation of viral E6 and E7 oncoproteins and down expression of viral proteins involved in viral particle assembly such as L1/L2 proteins are correlated with cancer progression[18, 19]. Epigenetic modification such as methylation of HPV genome is considered to be one factor to control the expression of viral genes during productive and transforming infections.

Differential methylation in HPV16 genome has been reported in cervical samples, during productive infection. HPV16 early promoter was found unmethylated in basal and intermediate cells at proximal E2 binding sites 2-4 (E2BS) but became highly methylated in superficial cells at the upper part of the epithelium. In latent HPV16 infection, viral long control region (LCR) including early promoter was highly methylated throughout the epithelium. In transforming HPV16 infected cells, the distal E2BS (E2BS1) and enhancer regions were found to be methylated, while the early promoter was unmethylated [20]. One study showed that HPV16 p670 late promoter was highly methylated in cervical carcinoma cases [21]. The low expression of viral early gene and lack of capsid L1/L2 proteins expression in undifferentiated basal cells could prevent activation of immune response to viral infection [22].

The methylation pattern in early promoter especially HPV16 E2BS has been widely studied in cervical cells and the methylation patterns were reported either progressive hypomethylation [23, 24] and progressive hypermethylation [21, 25, 26]. One study showed that high methylation of E2BS was correlated with episomal form in high grade cervical lesions, cervical carcinoma and multiple copies of integrated HPV genome [27]. L1 gene hypermethylation was correlated with severe cervical lesions and cervical carcinoma [28-30]. We have previously reported the association between high methylation of HPV16 L1 gene especially at CpG site 5600 and 5609 and high grade cervical lesions and cervical carcinoma [31]. However, there are limited studies of HPV16 genome methylation in anal cells especially in Asian countries. This study aimed to detect methylation pattern of HPV16 in early promoter and L1 regions in anal cells obtained from Thai MSM, analyzed by quantitative pyrosequencing assay. The pattern of methylation revealed the potential of using methylation at specific site as biomarker for screening of high-grade lesions of anal squamous intraepithelial cells.

## Materials and methods

### Clinical samples and cell lines

178 HPV16 positive DNA samples extracted from anal cells were collected from men who having sex with men (MSM) at the Thai Red Cross AIDS Research Centre (TRC-ARC) Bangkok, Thailand. There were 134 HIV positive cases and 44 HIV negative cases. Percentage of CD4+ results were obtained only from HIV positive men. DNA extracted from human cervical cancer cell lines containing of integrated HPV 16; CaSki, (CRL-1550 Lot No.3794357) containing approximately 500-600 copies per cell) and SiHa (HTB-35 Lot No.4031219) containing 1-2 copies per cell were used as positive control for amplification and pyrosequencing. This study has been approved (COA No. 053/2016) by the Institutional Review Board of the Faculty of Medicine, Chulalongkorn University.

### Methylation analysis by Pyrosequencing

The extracted DNA (100-1000 ng) from anal cells was bisulfite conversion by using EZ kit Gold Bisulfite Conversion Kit (Zymo Research) according to the manufacturer’s instruction. The sequences of forward, reverse and sequencing primers for early promoter CpG positions 31, 37, 43, 52 and 58, 3’L1 CpG positions 7136 and 7145 and 5’L1 CpG positions 5600, 5606, 5609 and 5615 were shown in Table 1.

**Table 1.**
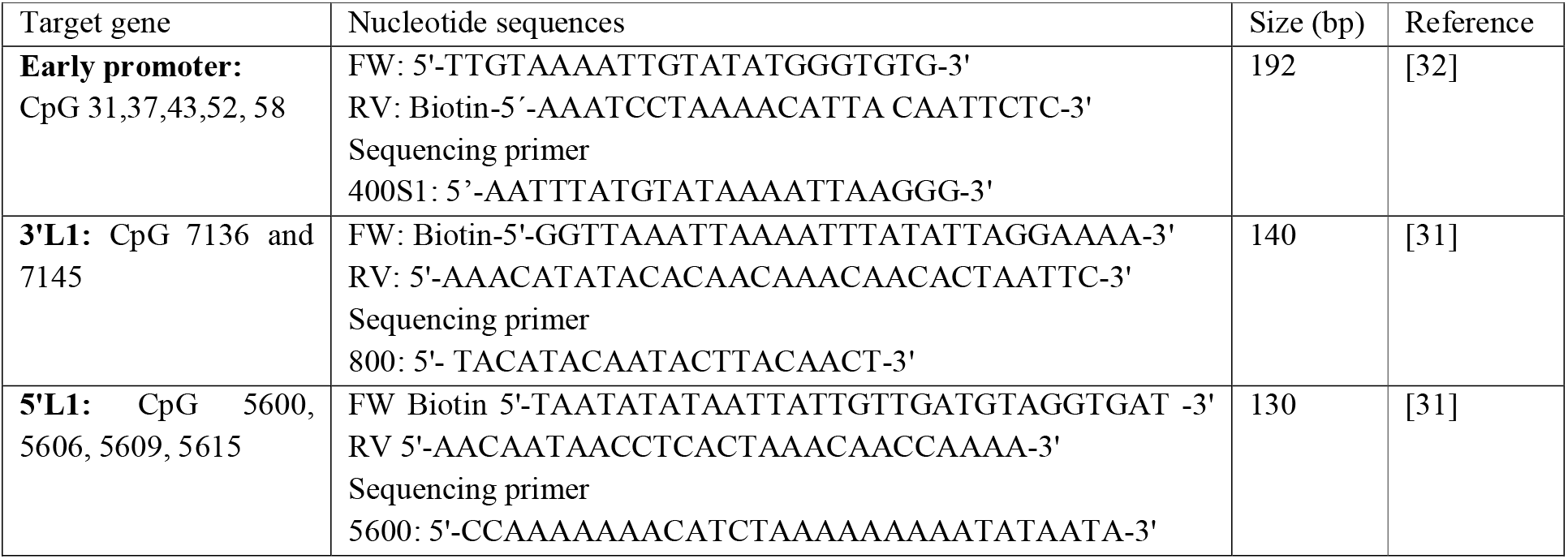
The sequences of forward and reverse primers used in this study

PCR amplification protocol was as follows: 13.6 μL DNase/RNase free water, 1X PCR buffer, 2.5mM MgCl_2_, 250μM dNTP, 12.5pM of each forward and reverse primers, 1 Unit DNA polymerase (HotStart HiFidelity Polymerase, Affymetrix, USA). The PCR conditions were started with initial denaturing at 95°C for 10 minutes, followed by 50 cycles of 95°C for 30 seconds, 55°C for 1 minute and 72°C for 1 minute and a cycle of final extension at 72°C for 10 minutes. The PCR products were detected by 1.5% agarose gel electrophoresis. Prior to pyrosequencing, 20μL of amplified products labeled with biotin were mixed with beads, washed, denatured, mixed with 0.4μM of sequencing primers and loaded into the PyroMark™ Q96 machine (Qiagen, Germany).

### Statistical analysis

The Kruskal-Wallis test was used to analyze the differences of the mean methylation values among groups of specimens. Fisher exact test was used to examine the significant differences of propotion of sample with methylation >20% between normal/AIN1 and AIN2-3. Pearson’s correlation coefficient (r) was used to analyse the association between percentage of methylation and CD4+. *P*-value less than 0.05 was considered statistically significant difference.

## Results

### Methylation levels of HPV16 early promoter and L1 gene in cervical cancer cell lines

Of 178 HPV16 positive samples including 134 HIV positive and 44 HIV negative samples, mean age was 31.23 years. Histology results were obtained from 123 samples classified as 16 normal, 67 AIN1, 12 AIN2 and 28 AIN3. Methylation patterns in HPV16 early promoter and two regions within HPV16 L1 gene were analyzed. The methylation levels of early promoter comprising 5 CpGs (31, 37, 43, 52 and 58) including proximal E2 binding sites (E2BSs) and Sp1 binding site of CaSki were 70%, 60.5%, 73.5%, 66.5%, 77% and of SiHa cells were 0-1% in all 5 CpGs, respectively. Methylation levels of L1 gene (5600, 5606, 5609, 5615, 7136 and 7145) were 84%, 59%, 76%, 65%, 69% and 67% for CaSki and 95%, 96%, 80%, 80%, 69% and 76% for SiHa, respectively, as shown in Fig 1. High methylation of L1 gene was detected in both cervical cell lines regardless of copy number of integrated HPV16.

**Fig 1.**
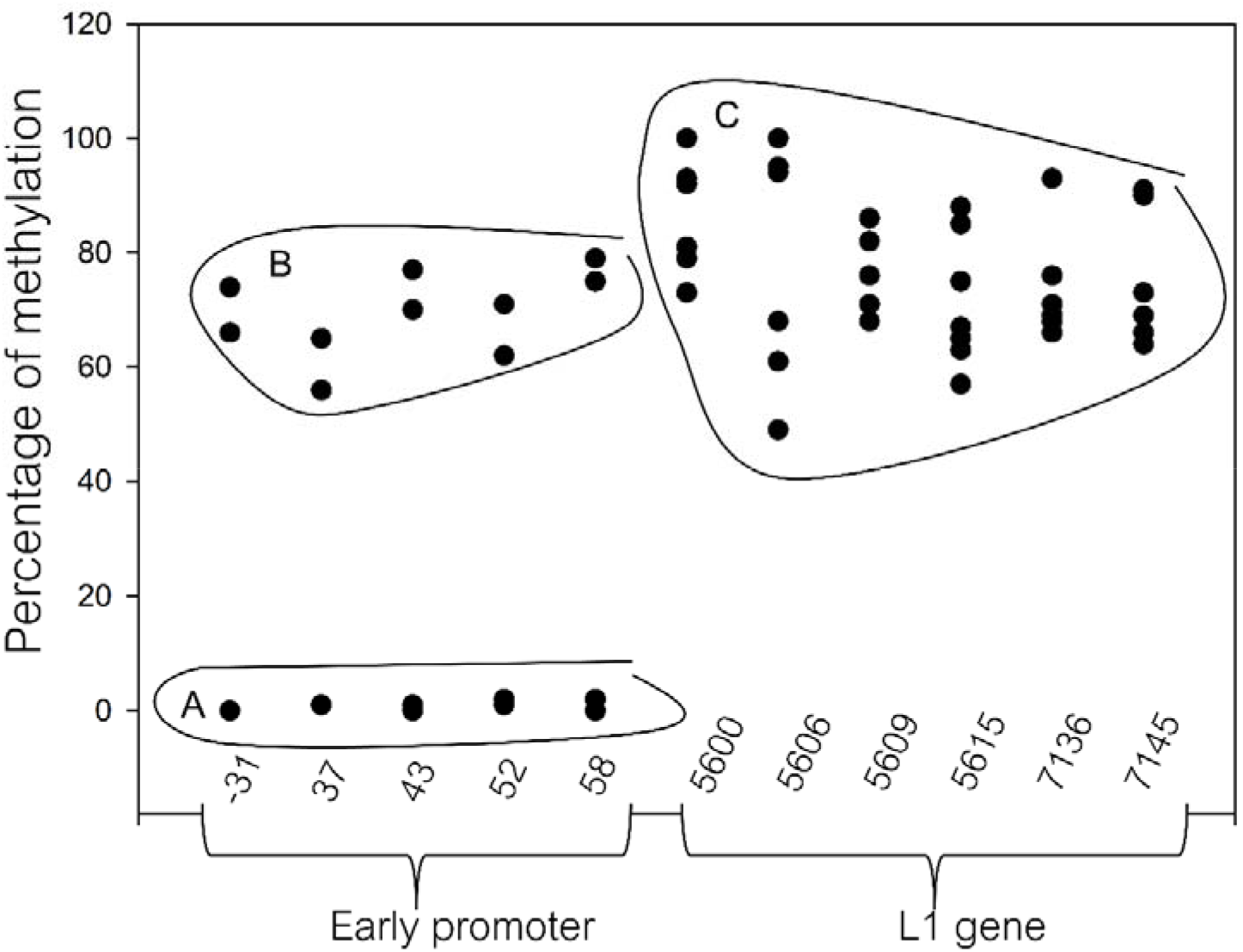
A scatter plot of methylation percentage of 11 CpGs located within early promoter (positions 31, 37, 43, 52 and 58), 3’L1 (positions 7136 and 7145), and 5’L1 regions (positions 5600, 5606, 5609 and 5615) of Caski and SiHa cell lines. A and B were methylation levels obtained from SiHa and CaSki, respectively. C was methylation level of L1 gene obtained from both SiHa and Caski.

### Methylation levels of HPV16 early promoter and L1 gene in anal cells

The methylation levels of early promoter in normal, AIN1 and AIN2-3 were shown in Fig 2. The methylation level was <10% in all normal anal cells, while intermediate methylation level (20-40%) was found in some of AIN1 and AIN2-3. The majority of AIN1 and AIN2-3 showed low methylation level in early promoter (<10%). For L1 gene, low methylation level was found in all detected normal anal cells (<20%). There was slightly increased in L1 gene methylation from normal to AIN2-3, especially at CpG5600 and 5609 which methylation levels were found in range 20-60% which were higher than other CpGs (5606, 5615, 7136 and 7145) (Fig 3). Intermediate to high methylation levels (20-60%) of CpG5600 was found in 0% (0/16) normal, 10.5% (7/67) AIN2-3 was found in CpG5609. There was significant difference of samples number with methylation percentage >20% between normal/AIN1 (8.4%; 7/83) and AIN2-3 (27.5%; 11/40) in CpG5600 (Table 2).

**Fig 2.**
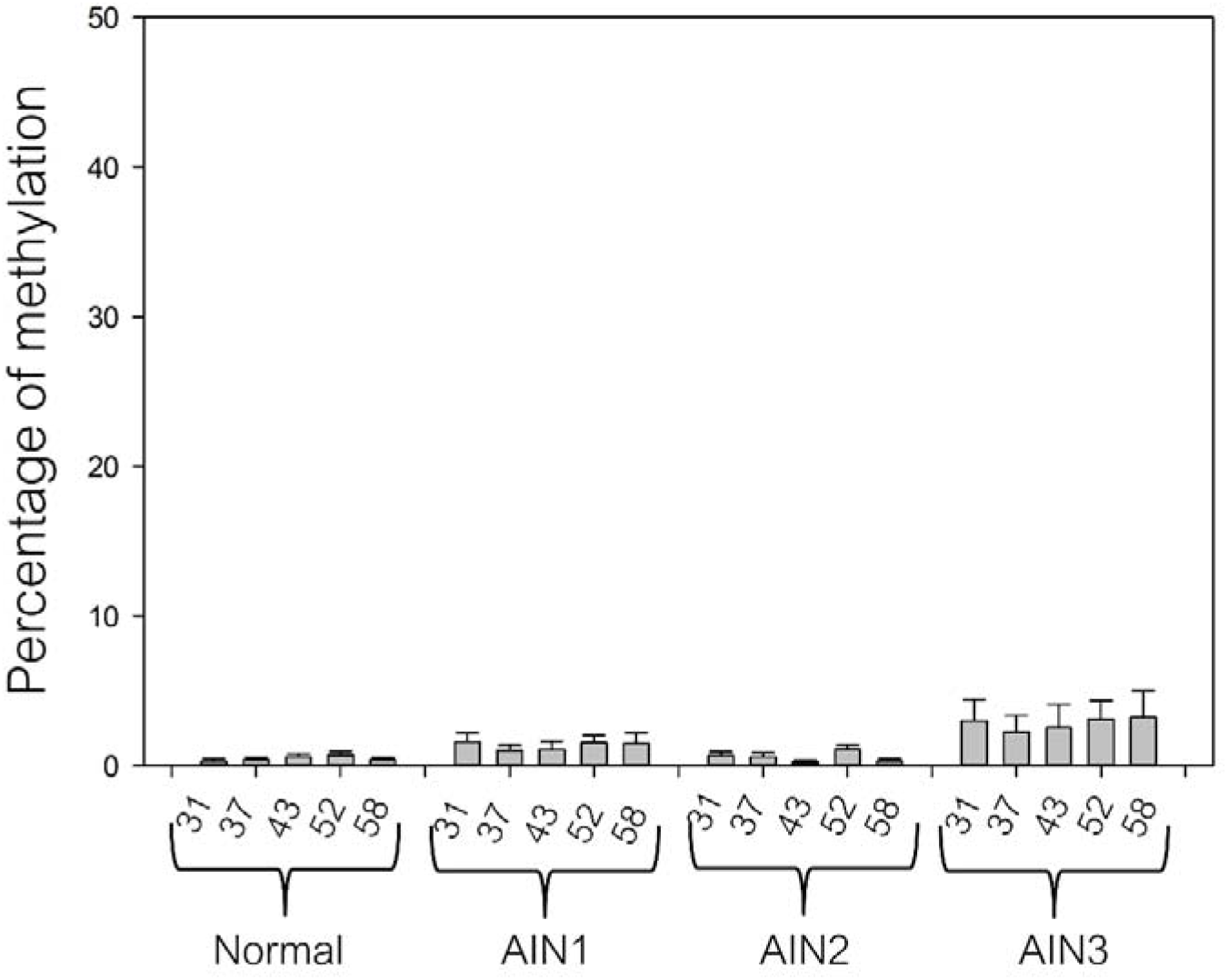
A bar graph of methylation percentage of early promoter in anal samples stratified by histology as normal, AIN1, AIN2 and AIN3 (mean±SE).

**Fig 3.**
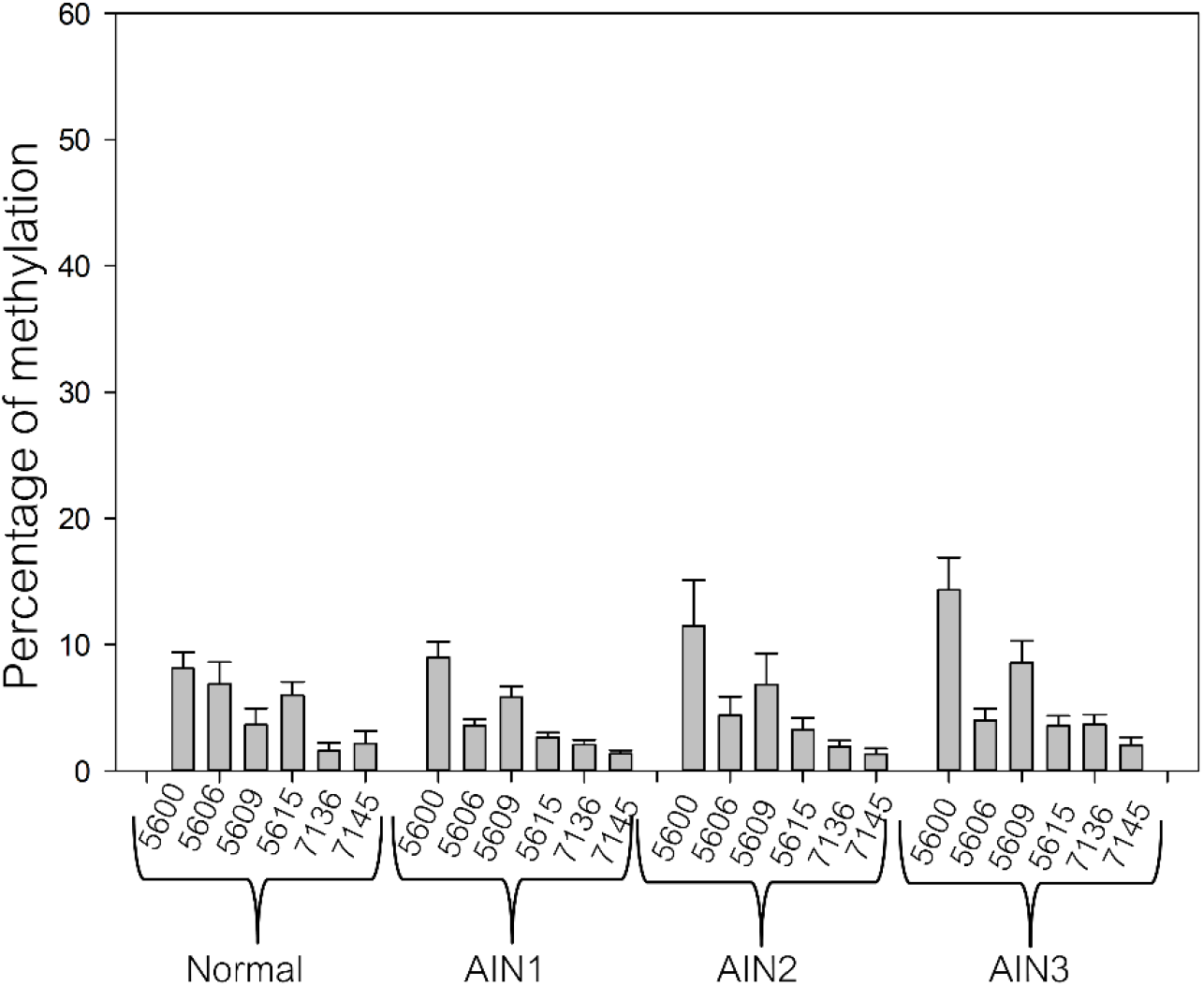
A bar graph of methylation percentage of L1 gene in anal samples stratified by histology as as normal, AIN1, AIN2 and AIN3 (mean±SE).

**Table 2.** Subject demographic and methylation status of HPV16 early promoter and L1 genes in anal cells with various grades of lesions. The *p*-values were calculated using Kruskal-Wallis test and Fisher Exact test. * indicated significant difference.

| Group | Normal | AIN1 | AIN2 | AIN3 | *p*-value |
| --- | --- | --- | --- | --- | --- |
| No. (123) | 16 | 67 | 12 | 28 |  |
| Age (years): |  |  |  |  |  |
| Mean | 27.93 | 31.06 | 31.3 | 32.52 | >0.05 |
| SD | 6.82 | 6.66 | 5.66 | 6.27 |  |
| Range | 22-47 | 19-50 | 25-42 | 22-52 |  |
| %Methylation level: |  |  |  |  |  |
| CpG 31 |  |  |  |  |  |
| Mean | 0.300 | 1.597 | 0.667 | 3.00 | >0.05 |
| SD | 0.483 | 4.568 | 0.888 | 7.299 |  |
| Range | 0-1 | 0-34 | 0-2 | 0-28 |  |
| % samples >20% | 0% | 1.5% | 0% | 7.1% | >0.05 |
| % samples <20% | 100% | 98.5% | 100% | 92.9% |  |
| CpG 37 |  |  |  |  |  |
| Mean | 0.400 | 1.016 | 0.583 | 2.269 | >0.05 |
| SD | 0.516 | 3.011 | 0.900 | 5.625 |  |
| Range | 0-1 | 0-23 | 0-3 | 0-23 |  |
| % samples >20% | 0% | 1.5% | 0% | 7.1% | >0.05 |
| % samples <20% | 100% | 98.5% | 100% | 92.9% |  |
| CpG 43 |  |  |  |  |  |
| Mean | 0.600 | 1.065 | 0.250 | 2.538 | >0.05 |
| SD | 0.699 | 4.745 | 0.452 | 7.824 |  |
| Range | 0-2 | 0-37 | 0-1 | 0-31 |  |
| % samples >20% | 0% | 1.5% | 0% | 7.1% | >0.05 |
| % samples <20% | 100% | 98.5% | 100% | 92.9% |  |
| CpG 52 |  |  |  |  |  |
| Mean | 0.700 | 1.532 | 1.083 | 3.115 | >0.05 |
| SD | 0.823 | 3.745 | 1.084 | 6.364 |  |
| Range | 0-2 | 0-28 | 0-3 | 0-26 |  |
| % samples >20% | 0% | 1.5% | 0% | 7.1% | >0.05 |
| % samples <20% | 100% | 98.5% | 100% | 92.9% |  |
| CpG 58 |  |  |  |  |  |
| Mean | 0.400 | 1.500 | 0.333 | 3.231 | >0.05 |
| SD | 0.516 | 5.331 | 0.492 | 8.878 |  |
| Range | 0-1 | 0-41 | 0-1 | 0-35 |  |
| % samples >20% | 0% | 1.5% | 0% | 7% | >0.05 |
| % samples <20% | 100% | 98.5% | 100% | 93% |  |
| CpG 5600 |  |  |  |  |  |
| Mean | 8.111 | 9.00 | 11.500 | 14.346 | >0.05 |
| SD | 3.887 | 8.976 | 12.494 | 13.069 |  |
| Range | 0-13 | 0-50 | 0-40 | 0-57 |  |
| % samples >20% | 0% | 10.5% | 25% | 28.6% | <0.05* |
| % samples <20% | 100% | 89.5% | 75% | 71.4% |  |
| CpG 5606 |  |  |  |  |  |
| Mean | 6.889 | 3.593 | 4.417 | 4.000 | >0.05 |
| SD | 5.183 | 3.616 | 5.017 | 4.699 |  |
| Range | 0-15 | 0-16 | 0-14 | 0-23 |  |
| % samples >20% | 22% | 0% | 0% | 3.6% | >0.05 |
| % samples <20% | 78% | 100% | 100% | 96.4% |  |
| CpG 5609 |  |  |  |  |  |
| Mean | 3.667 | 5.833 | 6.833 | 8.577 | >0.05 |
| SD | 3.841 | 6.279 | 8.537 | 8.732 |  |
| Range | 0-10 | 0-37 | 0-24 | 0-35 |  |
| % samples >20% | 0% | 3% | 16.7% | 7.1% | >0.05 |
| % samples <20% | 100% | 97% | 83.3% | 92.9% |  |
| CpG 5615 |  |  |  |  |  |
| Mean | 6.00 | 2.648 | 3.250 | 3.577 | >0.05 |
| SD | 3.12 | 2.789 | 3.251 | 4.032 |  |
| Range | 2-12 | 0-13 | 0-9 | 0-20 |  |
| % samples >20% | 0% | 0% | 0% | 3.6% | >0.05 |
| % samples <20% | 100% | 100% | 100% | 96.4% |  |
| CpG 7136 |  |  |  |  |  |
| Mean | 1.583 | 2.063 | 1.917 | 3.680 | >0.05 |
| SD | 2.193 | 2.945 | 1.621 | 3.945 |  |
| Range | 0-8 | 0-17 | 0-5 | 0-16 |  |
| % samples >20% | 0% | 0% | 0% | 8% | >0.05 |
| % samples <20% | 100% | 100% | 100% | 92% |  |
| CpG 7145 |  |  |  |  |  |
| Mean | 2.167 | 1.365 | 1.333 | 2.040 | >0.05 |
| SD | 3.460 | 2.180 | 1.557 | 3.089 |  |
| Range | 0-10 | 0-12 | 0-5 | 0-12 |  |
| % samples >20% | 0% | 0% | 0% | 8% | >0.05 |
| % samples <20% | 100% | 100% | 100% | 92% |  |

### Correlation between CD4+ percentage with HPV16 gene methylation

There was no statistically differences in mean of HPV16 methylation percentage between HIV negative and HIV positive cases (Table 3). Interestingly, HPV16 L1 gene high methylation was moderately correlated with low percentage of CD4+ in AIN3 HIV positive cases, especially at CpGs 5600, 5609, 5615 and 7145 (R=0.4692-0.5412) (Fig 4).

**Table 3.** Mean of HPV16 gene methylation of each CpGs, HIV status and percentage of CD4+

| Group | Normal (16) |  | AIN1(67) |  | AIN2(12) |  | AIN3(28) |  | p-value |
| --- | --- | --- | --- | --- | --- | --- | --- | --- | --- |
| HIV | negative | positive | negative | positive | negative | positive | negative | positive |  |
| No. (123) | 4 | 12 | 18 | 49 | 2 | 10 | 1 | 27 |  |
| Mean%CD4+ | NA | 19.5% | NA | 16.83% | NA | 19.2% | NA | 16.93% | >0.05 |
| Mean cell count |  | 389 |  | 315 |  | 359 |  | 326 |  |
| % Methylation |  |  |  |  |  |  |  |  |  |
| CpG 31 | 0% | 0.43% | 2.88% | 1.11% | 1.5% | 0.5% | 0% | 3.12% | >0.05 |
| CpG 37 | 0% | 0.57% | 2.0% | 0.64% | 2.0% | 0.3% | 0% | 2.36% | >0.05 |
| CpG 43 | 0% | 0.86% | 2.65% | 0.47% | 0% | 0.3% | 0% | 2.64% | >0.05 |
| CpG 52 | 0% | 1% | 2.47% | 1.18% | 2.0% | 0.9% | 0% | 3.24% | >0.05 |
| CpG 58 | 0% | 0.57% | 3.1% | 0.91% | 0.5% | 0.3% | 0% | 3.36% | >0.05 |
| CpG 5600 | 10.3% | 7% | 12.33% | 7.72% | 5.5% | 12.7% | 14% | 14.36% | >0.05 |
| CpG 5606 | 6.67% | 7% | 3.73% | 3.54% | 0% | 5.3% | 6% | 3.92% | >0.05 |
| CpG 5609 | 7% | 2% | 8.07% | 4.98% | 1.0% | 8.0% | 10% | 8.52% | >0.05 |
| CpG 5615 | 4.67% | 6.67% | 3.13% | 2.46% | 0% | 3.9% | 5% | 3.52% | >0.05 |
| CpG 7136 | 3% | 1.11% | 2.5% | 1.92% | 0.5% | 2.2% | 0% | 3.83% | >0.05 |
| CpG 7145 | 3.67% | 1.67% | 1.69% | 1.26% | 1.0% | 1.4% | 0% | 2.13% | >0.05 |

NA: not applicable

**Fig 4.**
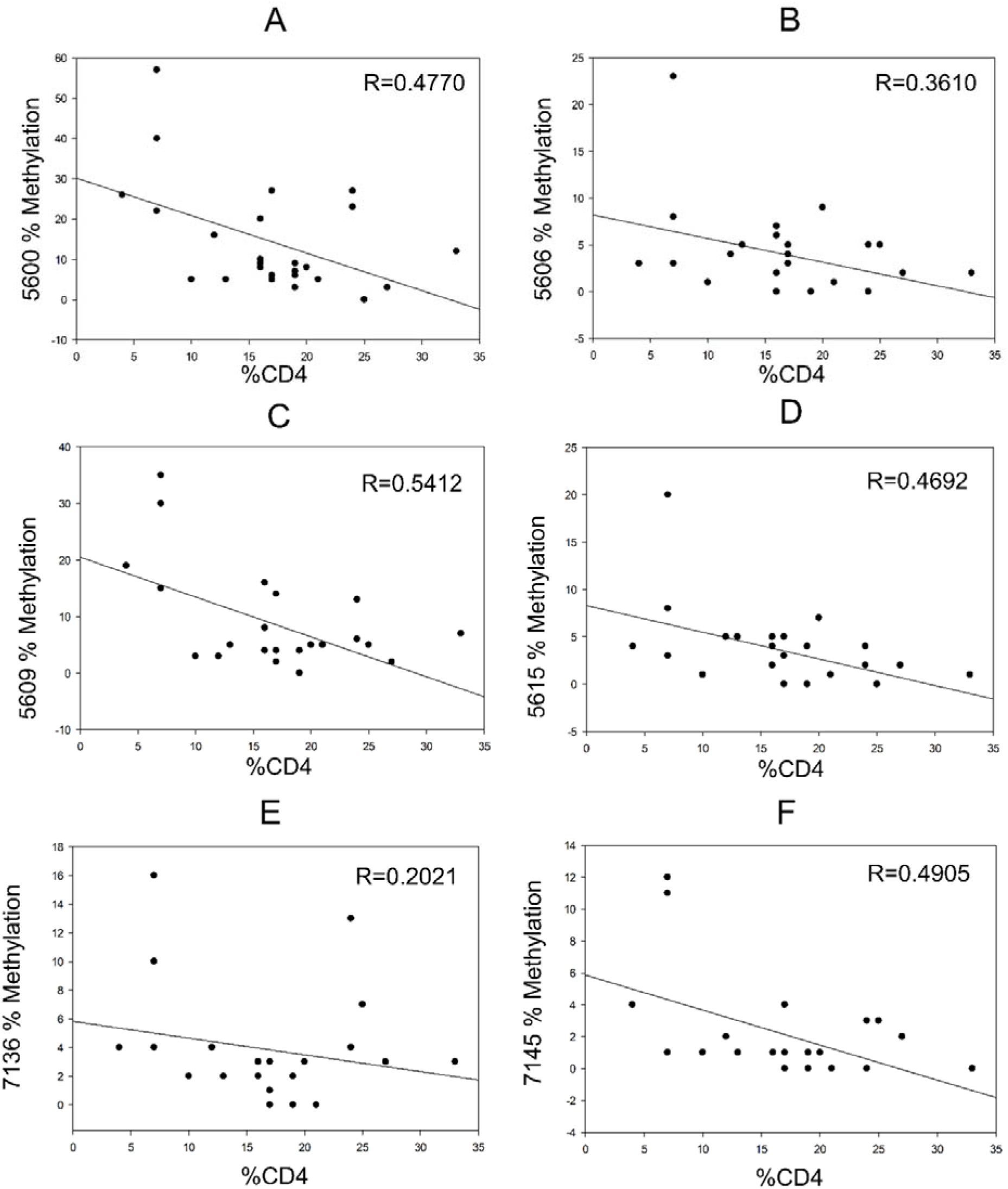
Correlation between percentage of HPV16 L1 gene methylation and number percentage of CD4+ in AIN3 cases.

## Discussion

In the present study, the methylation pattern of HPV16 genome including early promoter region and L1 gene in anal cells was studied using pyrosequencing assay. The quantitative methylation analysis was first analyzed in CaSki and SiHa cell lines that methylation levels were consistent with previous reports [29, 32–35]. HPV16 and 18 L1 gene hypermethylation was reported previously in cervical carcinoma, vulva intraepithelial neoplasia (VIN), oral carcinoma and penile carcinoma [29, 30, 36–41]. Hypermethylation in the L1 gene was found to be correlated with integration form of HPV16 [38, 42]. The present study found high methylation of HPV16 L1 gene in some of AIN2/3 samples compared to normal anal cells. We also found that CpG sites 5600 and 5609 showed higher methylation (>20%) compared to the other CpG sites in L1 region (5606, 5615, 7136 and 7145). Previous reports in cervical cells showed that CpG sites 5600 and 5609 were the best sites for separation of normal cervical cells and high grade dysplasia [29, 31, 43]. It can be implicit that methylation patterns of HPV16 L1 gene in anal cells were similar to cervical cells and might be used for separation of normal cells and HPV related severly abnormal cells.

Methylation patterns of HPV16 early promoter have been widely studied in cervical cells, nevertheless, controversial results were found. Some studies reported hypomethylation of HPV16 early promoter in cervical carcinoma or so-called progressive hypomethylation [23, 24]. Other studies reported hypermethylation of early promoter in cervical carcinoma or progressive hypermethylation [21, 25, 26, 30, 44]. The physical state and copy number of integrated HPV16 genomes were the main reasons of these methylation differences as shown in CaSki and SiHa cells. Episomal form of HPV16 genome found in high grade cervical lesions and cervical carcinoma displayed high methylation levels at E2 binding sites in early promoter compared with single integrated HPV16 genome [27]. One study showed high methylation of early promoter in high grade anal cells [45]. We can speculate that anal samples with intermediate to high methylation levels (>20%) in the present study may be at risk to progress quickly compared to those with low methylation (<20%).

It has been reported that HIV infected patients were susceptible for HPV infection. One study reported that in HIV infected women with CD4+ less than 200 cells/mm3 have 59.3% of high risk-HPV infections and correlated with increasing severity of cervical lesions [46]. The study in oral samples reported that low CD4+ count (<200 cells/mm3) increased risk for oral HPV infection in HIV infected patients [47]. In the present study, we could not get anal carcinoma samples due to very low incidence of anal cancer. Thus, in order to employ methylation of specific CpG sites for screening of HPV related cancer, a large sample size including anal carcinoma samples should be further completed and evaluated.

## Conclusions

The methylation patterns of HPV16 genome in anal intraepithelial neoplasia were similar to those of cervical abnormal cells. Hypermethylation in HPV16 L1 gene especially at CpG 5600 and 5609 found in AIN2/3 might be used for prediction of HPV related abnormal anal cells. Moreover, high HPV16 L1 gene methylation together with low CD4+ cell count in HIV infected patients may be used as the biomarker for rapid progression to more severely lesions than those with low methylation and high CD4+ cell count.

## Acknowledgments

This work was supported by Research Grant for New Scholar Ratchadaphiseksomphot Endowment Fund, Chulalongkorn University.

## Notes

### Competing Interest Statement

The authors have declared no competing interest.

## References

1. Flejou JF. An update on anal neoplasia. Histopathology. 2015;66(1): 147–60. doi: 10.1111/his.12574. PubMed PMID: 25283345.

2. Wang CJ, Sparano J, Palefsky JM. Human Immunodeficiency Virus/AIDS, Human Papillomavirus, and Anal Cancer. Surg Oncol Clin N Am. 2017;26(1):17–31. Epub 2016/11/28. doi: 10.1016/j.soc.2016.07.010. PubMed PMID: 27889034; PubMed Central PMCID: PMCPMC5331942.

3. De Vuyst H, Clifford GM, Nascimento MC, Madeleine MM, Franceschi S. Prevalence and type distribution of human papillomavirus in carcinoma and intraepithelial neoplasia of the vulva, vagina and anus: a meta-analysis. Int J Cancer. 2009; 124(7): 1626–36. Epub 2008/12/31. doi: 10.1002/ijc.24116. PubMed PMID: 19115209.

4. Serrano B, de Sanjose S, Tous S, Quiros B, Munoz N, Bosch X, et al. Human papillomavirus genotype attribution for HPVs 6, 11, 16, 18, 31, 33, 45, 52 and 58 in female anogenital lesions. Eur J Cancer. 2015. Epub 2015/07/01. doi: S0959-8049(15)00491-8 [pii]10.1016/j.ejca.2015.06.001. PubMed PMID: 26121913.

5. Daling JR, Madeleine MM, Johnson LG, Schwartz SM, Shera KA, Wurscher MA, et al. Human papillomavirus, smoking, and sexual practices in the etiology of anal cancer. Cancer. 2004;101(2):270–80. Epub 2004/07/09. doi: 10.1002/cncr.20365. PubMed PMID: 15241823.

6. Machalek DA, Poynten M, Jin F, Fairley CK, Farnsworth A, Garland SM, et al. Anal human papillomavirus infection and associated neoplastic lesions in men who have sex with men: a systematic review and meta-analysis. Lancet Oncol. 2012;13(5):487–500. Epub 2012/03/27. doi: S1470-2045(12)70080-3 [pii]10.1016/S1470-2045(12)70080-3. PubMed PMID: 22445259.

7. Li X, Li M, Yang Y, Zhong X, Feng B, Xin H, et al. Anal HPV/HIV co-infection among Men Who Have Sex with Men: a cross-sectional survey from three cities in China. Scientific reports. 2016;6:21368. doi: 10.1038/srep21368. PubMed PMID: 26892938; PubMed Central PMCID: PMC4759533.

8. Phanuphak N, Teeratakulpisarn N, Pankam T, Kerr SJ, Barisri J, Deesua A, et al. Anal human papillomavirus infection among Thai men who have sex with men with and without HIV infection: prevalence, incidence, and persistence. J Acquir Immune Defic Syndr. 2013;63(4):472–9. doi: 10.1097/QAI.0b013e3182918a5a. PubMed PMID: 23514956; PubMed Central PMCID: PMC3700660.

9. Supindham T, Chariyalertsak S, Utaipat U, Miura T, Ruanpeng D, Chotirosniramit N, et al. High Prevalence and Genotype Diversity of Anal HPV Infection among MSM in Northern Thailand. PloS one. 2015; 10(5):e0124499. doi: 10.1371/journal.pone.0124499. PubMed PMID: 25932915; PubMed Central PMCID: PMC4416722.

10. Cho HW, So KA, Lee JK, Hong JH. Type-specific persistence or regression of human papillomavirus genotypes in women with cervical intraepithelial neoplasia 1: A prospective cohort study. Obstetrics & gynecology science. 2015;58(1):40–5. doi: 10.5468/ogs.2015.58.1.40. PubMed PMID: 25629017; PubMed Central PMCID: PMC4303751.

11. Trimble CL, Piantadosi S, Gravitt P, Ronnett B, Pizer E, Elko A, et al. Spontaneous regression of high-grade cervical dysplasia: effects of human papillomavirus type and HLA phenotype. Clin Cancer Res. 2005;11(13):4717–23. Epub 2005/07/08. doi: 11/13/4717 [pii] 10.1158/1078-0432.CCR-04-2599. PubMed PMID: 16000566; PubMed Central PMCID: PMC3132609.

12. Schlecht NF, Platt RW, Duarte-Franco E, Costa MC, Sobrinho JP, Prado JC, et al. Human papillomavirus infection and time to progression and regression of cervical intraepithelial neoplasia. J Natl Cancer Inst. 2003;95(17): 1336–43. PubMed PMID: 12953088.

13. Dona MG, Vescio MF, Latini A, Giglio A, Moretto D, Frasca M, et al. Anal human papillomavirus in HIV-uninfected men who have sex with men: incidence and clearance rates, duration of infection, and risk factors. Clinical microbiology and infection: the official publication of the European Society of Clinical Microbiology and Infectious Diseases. 2016;22(12): 1004 e1–e7. doi: 10.1016/j.cmi.2016.08.011. PubMed PMID: 27585942.

14. Munger K, Baldwin A, Edwards KM, Hayakawa H, Nguyen CL, Owens M, et al. Mechanisms of human papillomavirus-induced oncogenesis. J Virol. 2004;78(21):11451–60. PubMed PMID: 15479788.

15. Munger K, Basile JR, Duensing S, Eichten A, Gonzalez SL, Grace M, et al. Biological activities and molecular targets of the human papillomavirus E7 oncoprotein. Oncogene. 2001;20(54):7888–98. PubMed PMID: 11753671.

16. Chen HC, Schiffman M, Lin CY, Pan MH, You SL, Chuang LC, et al. Persistence of type-specific human papillomavirus infection and increased long-term risk of cervical cancer. J Natl Cancer Inst. 2011;103(18):1387–96. doi: 10.1093/jnci/djr283. PubMed PMID: 21900119; PubMed Central PMCID: PMC3176778.

17. Moscicki AB, Shiboski S, Hills NK, Powell KJ, Jay N, Hanson EN, et al. Regression of low-grade squamous intra-epithelial lesions in young women. Lancet. 2004;364(9446): 1678–83. Epub 2004/11/09. doi: S0140673604173546 [pii] 10.1016/S0140-6736(04)17354-6. PubMed PMID: 15530628.

18. Xiao W, Bian ML, Ma L, Liu J, Chen Y. [Detection of human papillomavirus L1 capsid protein expression in liquid-based cytology samples with abnormal cytology.]. Zhonghua Fu Chan Ke Za Zhi. 2009;44(12):887–91. Epub 2010/03/03. PubMed PMID: 20193413.

19. Origoni M, Cristoforoni P, Carminati G, Stefani C, Costa S, Sandri MT, et al. *E6/E1* mRNA testing for human papilloma virus-induced high-grade cervical intraepithelial disease (CIN2/CIN3): a promising perspective. Ecancermedicalscience. 2015;9:533. Epub 2015/05/28. doi: 10.3332/ecancer.2015.533 can-9-533 [pii]. PubMed PMID: 26015802; PubMed Central PMCID: PMC4435751.

20. Vinokurova S, von Knebel Doeberitz M. Differential methylation of the HPV16 upstream regulatory region during epithelial differentiation and neoplastic transformation. PloS one. 2011;6(9):e24451. PubMed PMID: 21915330.

21. Baedyananda F, Chaiwongkot A, Bhattarakosol P. Elevated HPV16 E1 Expression Is Associated with Cervical Cancer Progression. Intervirology. 2017;60(5): 171–80. doi: 10.1159/000487048. PubMed PMID: 29495005.

22. Burley M, Roberts S, Parish JL. Epigenetic regulation of human papillomavirus transcription in the productive virus life cycle. Seminars in Immunopathology. 2020;42(2): 159–71. doi: 10.1007/s00281-019-00773-0.

23. Badal V, Chuang LS, Tan EH, Badal S, Villa LL, Wheeler CM, et al. CpG methylation of human papillomavirus type 16 DNA in cervical cancer cell lines and in clinical specimens: genomic hypomethylation correlates with carcinogenic progression. J Virol. 2003;77(11):6227–34. Epub 2003/05/14. PubMed PMID: 12743279; PubMed Central PMCID: PMC 154984.

24. Piyathilake CJ, Macaluso M, Alvarez RD, Chen M, Badiga S, Edberg JC, et al. A higher degree of methylation of the HPV 16 E6 gene is associated with a lower likelihood of being diagnosed with cervical intraepithelial neoplasia. Cancer. 2011;117(5):957–63. Epub 2010/10/15. doi: 10.1002/cncr.25511. PubMed PMID: 20945322; PubMed Central PMCID: PMC3023831.

25. Bhattacharjee B, Sengupta S. CpG methylation of HPV 16 LCR at E2 binding site proximal to *P97* is associated with cervical cancer in presence of intact E2. Virology. 2006;354(2):280–5. PubMed PMID: 16905170.

26. Ding DC, Chiang MH, Lai HC, Hsiung CA, Hsieh CY, Chu TY. Methylation of the long control region of HPV16 is related to the severity of cervical neoplasia. Eur J Obstet Gynecol Reprod Biol. 2009;147(2):215–20. Epub 2009/10/13. doi: 10.1016/j.ejogrb.2009.08.023 S0301-2115(09)00507-7 [pii]. PubMed PMID: 19819061.

27. Chaiwongkot A, Vinokurova S, Pientong C, Ekalaksananan T, Kongyingyoes B, Kleebkaow P, et al. Differential methylation of E2 binding sites in episomal and integrated HPV 16 genomes in preinvasive and invasive cervical lesions. Int J Cancer. 2013;132(9):2087–94. Epub 2012/10/16. doi: 10.1002/ijc.27906. PubMed PMID: 23065631.

28. Oka N, Kajita M, Nishimura R, Ohbayashi C, Sudo T. L1 gene methylation in high-risk human papillomaviruses for the prognosis of cervical intraepithelial neoplasia. Int J Gynecol Cancer. 2013;23(2):235–43. Epub 2013/01/15. doi: 10.1097/IGC.0b013e31827da1f6. PubMed PMID: 23314283.

29. Bryant D, Tristram A, Liloglou T, Hibbitts S, Fiander A, Powell N. Quantitative measurement of Human Papillomavirus type 16 L1/L2 DNA methylation correlates with cervical disease grade. J Clin Virol. 2014;59(1):24–9. Epub 2013/11/26. doi: 10.1016/j.jcv.2013.10.029 S1386-6532(13)00477-0 [pii]. PubMed PMID: 24268385.

30. Fernandez AF, Rosales C, Lopez-Nieva P, Grana O, Ballestar E, Ropero S, et al. The dynamic DNA methylomes of double-stranded DNA viruses associated with human cancer. Genome Res. 2009;19(3):438–51. Epub 2009/02/12. doi: 10.1101/gr.083550.108gr.083550.108 [pii]. PubMed PMID: 19208682; PubMed Central PMCID: PMC2661803.

31. Chaiwongkot A, Niruthisard S, Kitkumthorn N, Bhattarakosol P. Quantitative methylation analysis of human papillomavirus 16L1 gene reveals potential biomarker for cervical cancer progression. Diagn Microbiol Infect Dis. 2017;89(4):265–70. doi: 10.1016/j.diagmicrobio.2017.08.010. PubMed PMID: 28985972.

32. Rajeevan MS, Swan DC, Duncan K, Lee DR, Limor JR, Unger ER. Quantitation of site-specific HPV 16 DNA methylation by pyrosequencing. J Virol Methods. 2006; 138(1-2):170–6. PubMed PMID: 17045346.

33. Kalantari M, Osann K, Calleja-Macias IE, Kim S, Yan B, Jordan S, et al. Methylation of human papillomavirus 16, 18, 31, and 45 L2 and L1 genes and the cellular DAPK gene: Considerations for use as biomarkers of the progression of cervical neoplasia. Virology. 2014;448:314–21. Epub 2013/12/10. doi: 10.1016/j.virol.2013.10.032 S0042-6822(13)00607-7 [pii]. PubMed PMID: 24314662.

34. Lorincz AT, Brentnall AR, Vasiljevic N, Scibior-Bentkowska D, Castanon A, Fiander A, et al. HPV16 L1 and L2 DNA methylation predicts high-grade cervical intraepithelial neoplasia in women with mildly abnormal cervical cytology. Int J Cancer. 2013;133(3):637–44. Epub 2013/01/22. doi: 10.1002/ijc.28050. PubMed PMID: 23335178; PubMed Central PMCID: PMC3708123.

35. Brandsma JL, Harigopal M, Kiviat NB, Sun Y, Deng Y, Zelterman D, et al. Methylation of Twelve CpGs in Human Papillomavirus Type 16 (HPV16) as an Informative Biomarker for the Triage of Women Positive for HPV16 Infection. Cancer prevention research. 2014;7(5):526–33. Epub 2014/02/22. doi: 10.1158/1940-6207.CAPR-13-0354 1940-6207.CAPR-13-0354 [pii]. PubMed PMID: 24556390.

36. Kalantari M, Villa LL, Calleja-Macias IE, Bernard HU. Human papillomavirus-16 and −18 in penile carcinomas: DNA methylation, chromosomal recombination and genomic variation. Int J Cancer. 2008; 123(8): 1832–40. Epub 2008/08/09. doi: 10.1002/ijc.23707. PubMed PMID: 18688866; PubMed Central PMCID: PMC2750853.

37. Balderas-Loaeza A, Anaya-Saavedra G, Ramirez-Amador VA, Guido-Jimenez MC, Kalantari M, Calleja-Macias IE, et al. Human papillomavirus-16 DNA methylation patterns support a causal association of the virus with oral squamous cell carcinomas. Int J Cancer. 2007;120(10):2165–9. Epub 2007/02/06. doi: 10.1002/ijc.22563. PubMed PMID: 17278110.

38. Bryant D, Onions T, Raybould R, Jones S, Tristram A, Hibbitts S, et al. Increased methylation of Human Papillomavirus type 16 DNA correlates with viral integration in Vulval Intraepithelial Neoplasia. J Clin Virol. 2014;61(3):393–9. Epub 2014/09/15. doi: 10.1016/j.jcv.2014.08.006 S1386-6532(14)00306-0 [pii]. PubMed PMID: 25218242.

39. Badal S, Badal V, Calleja-Macias IE, Kalantari M, Chuang LS, Li BF, et al. The human papillomavirus-18 genome is efficiently targeted by cellular DNA methylation. Virology. 2004;324(2):483–92. Epub 2004/06/23. doi: 10.1016/j.virol.2004.04.002 S0042682204002326 [pii]. PubMed PMID: 15207633.

40. Sun C, Reimers LL, Burk RD. Methylation of HPV16 genome CpG sites is associated with cervix precancer and cancer. Gynecol Oncol. 2011;121(1):59–63. Epub 2011/02/11. doi: S0090-8258(11)00046-1 [pii] 10.1016/j.ygyno.2011.01.013. PubMed PMID: 21306759; PubMed Central PMCID: PMC3062667.

41. Turan T, Kalantari M, Calleja-Macias IE, Cubie HA, Cuschieri K, Villa LL, et al. Methylation of the human papillomavirus-18 L1 gene: a biomarker of neoplastic progression? Virology. 2006;349(1): 175–83. Epub 2006/02/14. doi: S0042-6822(05)00858-5 [pii] 10.1016/j.virol.2005.12.033. PubMed PMID: 16472835.

42. Kalantari M, Chase DM, Tewari KS, Bernard HU. Recombination of human papillomavirus-16 and host DNA in exfoliated cervical cells: a pilot study of L1 gene methylation and chromosomal integration as biomarkers of carcinogenic progression. J Med Virol. 2010;82(2):311–20. PubMed PMID: 20029805.

43. Bryant D, Hibbitts S, Almonte M, Tristram A, Fiander A, Powell N. Human papillomavirus type 16 L1/L2 DNA methylation shows weak association with cervical disease grade in young women. J Clin Virol. 2015;66:66–71. Epub 2015/04/14. doi: 10.1016/j.jcv.2015.03.001 S1386-6532(15)00078-5 [pii]. PubMed PMID: 25866341.

44. Kalantari M, Calleja-Macias IE, Tewari D, Hagmar B, Lie K, Barrera-Saldana HA, et al. Conserved methylation patterns of human papillomavirus type 16 DNA in asymptomatic infection and cervical neoplasia. J Virol. 2004;78(23): 12762–72. Epub 2004/11/16. doi: 78/23/12762 [pii] 10.1128/JVI.78.23.12762-12772.2004. PubMed PMID: 15542628; PubMed Central PMCID: PMC525027.

45. Wiley DJ, Huh J, Rao JY, Chang C, Goetz M, Poulter M, et al. Methylation of human papillomavirus genomes in cells of anal epithelia of HIV-infected men. J Acquir Immune Defic Syndr. 2005;39(2): 143–51. Epub 2005/05/21. doi: 00126334-200506010-00004 [pii]. PubMed PMID: 15905729.

46. Teixeira MF, Sabidó M, Leturiondo AL, de Oliveira Ferreira C, Torres KL, Benzaken AS. High risk human papillomavirus prevalence and genotype distribution among women infected with HIV in Manaus, Amazonas. Virology Journal. 2018;15(1):36. doi: 10.1186/s12985-018-0942-6.

47. Muller K, Kazimiroff J, Fatahzadeh M, Smith RV, Wiltz M, Polanco J, et al. Oral Human Papillomavirus Infection and Oral Lesions in HIV-Positive and HIV-Negative Dental Patients. The Journal of infectious diseases. 2015;212(5):760–8. Epub 02/13. doi: 10.1093/infdis/jiv080. PubMed PMID: 25681375.

